# Achieving single nucleotide sensitivity in direct hybridization genome imaging

**DOI:** 10.1101/2022.05.31.494230

**Authors:** Yanbo Wang, Wayne T. Cottle, Haobo Wang, Momcilo Gavrilov, Roger S. Zou, Scott Bailey, Taekjip Ha

## Abstract

Direct visualization of point mutations *in situ* can be informative for studying genetic diseases and nuclear biology. We describe a direct hybridization genome imaging method with single-nucleotide sensitivity, sgGOLDFISH, which leverages the high cleavage specificity of enhanced Cas9 combined with a single extended guide RNA to load a superhelicase and reveal probe binding sites through local denaturation. Using sgGOLDFISH, we identified base-editor-modified and unmodified progeroid fibroblasts from a heterogeneous population, validated the identification through progerin immunofluorescence, and demonstrated accurate sub-nuclear localization of point mutations.

## Main text

Single-nucleotide variation (SNV) is the most common type of mutation and is associated with many diseases^1^. Although sequencing approaches can detect SNVs, they do not report on spatial information. Fluorescence *in situ* hybridization (FISH) can reveal the three-dimensional location of genomic sites of interest through annealing of fluorescently labeled oligonucleotide probes to denatured chromosomal DNA, but generally cannot differentiate highly similar sequences. Advanced DNA and RNA FISH methods have been developed to visualize SNVs. However, SNV-sensitive RNA FISH requires the target RNA to be actively transcribing, thereby excluding nongenic regions and inactive or stochastically expressed genes^2-8^. SNV-sensitive DNA FISH utilizes *in situ* PCR or rolling circle amplification strategy to detect SNVs, but the visualized signals come from bulky amplification products instead of the target SNVs^9-11^. The vast majority of genome imaging has been performed through direct probe hybridization to target chromatin, and to date, direct hybridization DNA FISH with SNV sensitivity has not been realized. We developed single guide (sg) version of GOLDFISH (genome oligopaint via local denaturation FISH) to address this technical gap.

The original GOLDFISH method utilized multiple guide RNAs tiling a genomic region of interest in complex with Cas9 nickase (SpCas9 with H840A mutation^12^) to nick genomic DNA at multiple sites (Extended Data Fig. 1a)^13^. This allowed local denaturation of targeted genomic DNA by loading an engineered helicase, Rep-X, to the cleaved strands so that DNA downstream is unwound to expose binding sites for FISH probes. Because only several kilobases of DNA are unwound, GOLDFISH greatly reduces nonspecific binding of FISH probes to other genomic regions compared to conventional FISH that globally denatures genomic DNA. Leveraging the increased sequence-stringency of Cas9 cleavage compared to Cas9 binding, GOLDFISH also has superior signal-to-background ratios compared to methods that rely on Cas9 binding^13^.

The use of multiple cut sites in GOLDFISH enabled high efficiency labeling even if the cleavage efficiency at a single site is not very high, but it prevented SNV detection. We hypothesized that, instead of using multiple guide RNAs, GOLDFISH using a single guide RNA (hence called sgGOLDFISH, Extended Data Fig. 1b) may achieve SNV sensitivity if the Cas9 cleavage activity is optimized to be SNV-sensitive (Fig. 1a). If so, by rationally designing guide RNA and choosing an engineered Cas9 variant with higher cleavage specificity, sgGOLDFISH could preferentially label one of the two alleles even when the two alleles differ by only a single nucleotide (Fig. 1a). To test this concept, we assembled a Cas9 ribonucleoprotein (RNP) variant by combining the specificity-improving mutations in eSpCas9(1.1)^14^ with a 5’ extended guide RNA^15^ (hereinafter called eCas9 RNP, which cleaves both the target and nontarget strands, Extended Data Fig. 2a), and tested its cleavage activity *in vitro* (Extended Data Fig. 2b and 2c). We observed efficient cleavage for 4 out of 5 guide RNAs with 1 PAM-distal mismatch, but no cleavage activity for five guide RNAs containing 2 PAM-distal mismatches (Extended Data Fig. 2d). Such specificity was not observed with canonical guide RNA in complex with eCas9 (Extended Data Fig. 2e). These data indicate that adding an extra PAM-distal mismatch can drastically reduce the cleavage activity of eCas9 RNP.

**Fig. 1.**
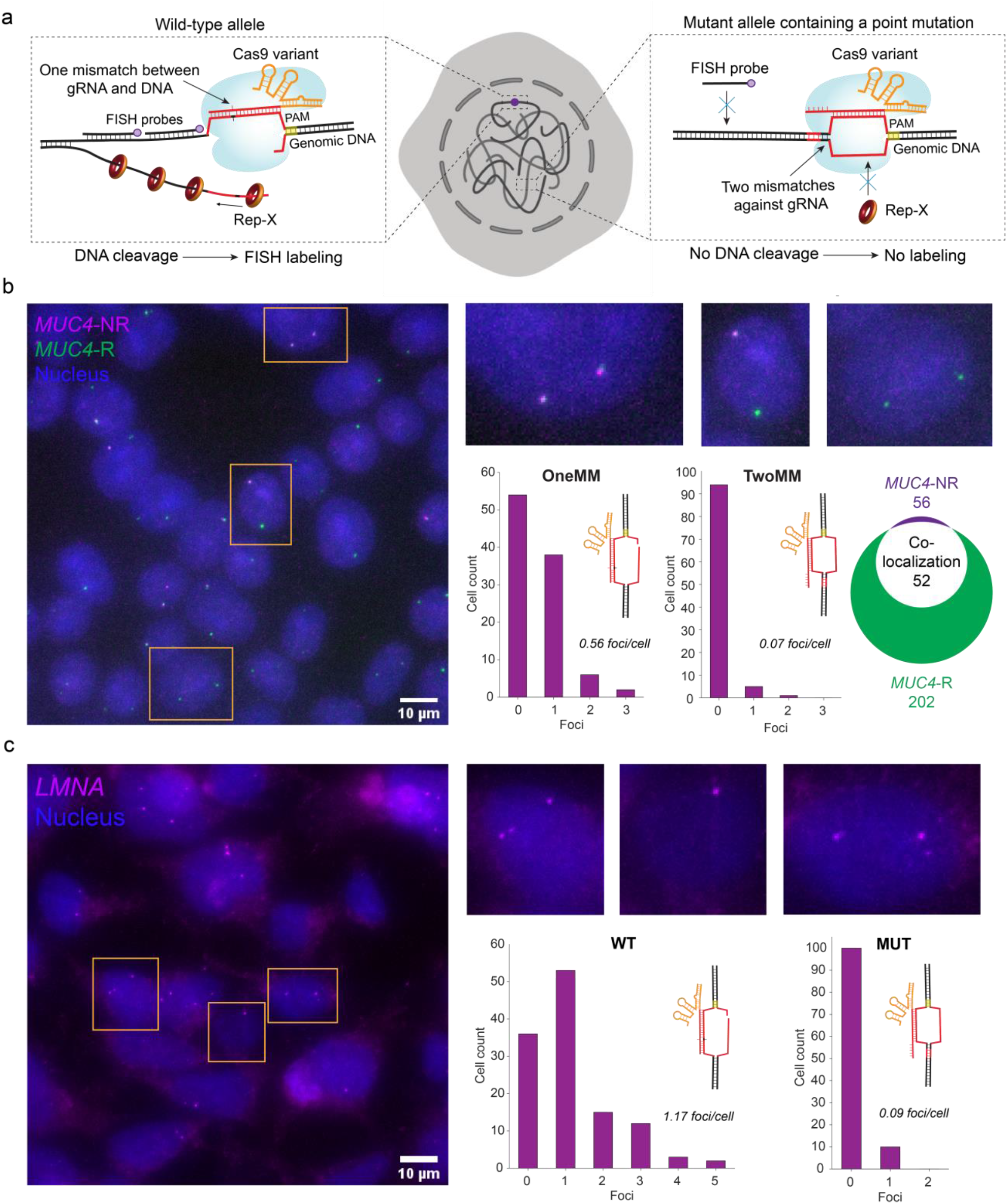
sgGOLDFISH. **a**, Schematic of SNV detection using sgGOLDFISH. Genomic DNA in red is homologous to guide RNA. **b**, A representative sgGOLDFISH image using gMUC4-OneMM (single cells outlined in orange are magnified on the upper–right corner), histograms of sgGOLDFISH *MUC4*-NR foci per cell using gMUC4-OneMM (n=78) or gMUC4-TwoMM (n=100), and quantification of co-localized *MUC4*-R and *MUC4*-NR foci. **c**, A representative sgGOLDFISH image using gLMNA-WT (single cells outlined in orange are magnified on the upper–right corner), and histograms of sgGOLDFISH *LMNA* foci using gLMNA-WT (n=121) or gLMNA-MUT (n=110).

Cleaving only the non-target strand by Cas9 nickase is sufficient for Rep-X unwinding the downstream genomic DNA and FISH probe hybridization in GOLDFISH^13^. We therefore created the eCas9(H840A) variant (hereinafter called eCas9 nickase)^12^, and measured its cleavage activity in fixed cells (note that in GOLDFISH, cleavage and subsequent steps are performed in fixed cells). The fraction of double-strand breaks (DSBs) at a target site in the cell population can be measured using a droplet digital PCR (ddPCR) assay^16^. ddPCR, however, is not sensitive to single-strand breaks (SSBs). Here we extended the ddPCR assay through an additional nicking step to make it SSB-sensitive (Extended Data Fig. 3-5, see Supplementary Note for detailed descriptions). Using this assay, we measured cleavage efficiency of eCas9 nickase in complex with a guide RNA that contains two mismatches against the *MUC4* gene (gMUC4-TwoMM), or a guide RNA with one mismatch (gMUC4-OneMM), and we observed < 5 % and ∼ 40 % DNA being cleaved, respectively (Extended Data Fig. 3b and 3c). These data demonstrate that an extra mismatch drastically reduced the cleavage efficiency of eCas9 nickase RNP in fixed cells so that a contrast of about an order of magnitude can be obtained.

We first tested sgGOLDFISH in proteinase-treated cells (HEK293T) using eCas9 nickase with gMUC4-OneMM or gMUC4-TwoMM alongside 23 Cy5-labeled FISH probes against a 1.5-kb non-repetitive region in the *MUC4* gene (*MUC4*-NR) adjacent to the target protospacer (Extended Data Fig. 6a and 6b). Another guide RNA (gMUC4-R) and a Cy3-labeled FISH probe were also designed against a repetitive region (*MUC4*-R) 19-kb from the *MUC4*-NR region to evaluate the specificity and sensitivity of sgGOLDFISH (Extended Data Fig. 6a). With gMUC4-OneMM, on average 0.56 *MUC4*-NR FISH foci per cell (foci/cell) were detected (Fig. 1b). Whereas with gMUC4-TwoMM, on average 0.07 foci/cell were detected (Fig. 1b). With concurrent use of gMUC4-R and gMUC4-OneMM, we found that 52 of 56 *MUC4*-NR foci colocalized with *MUC4-R* foci, indicating high labeling specificity of sgGOLDFISH (Fig. 1b). There were a total of 202 *MUC4*-R foci among which 52 showed colocalization with *MUC4*-NR, suggesting about 26% detection efficiency of sgGOLDFISH. We then performed sgGOLDFISH without proteinase treatment and observed 0.45 foci/cell for gMUC4-OneMM and 0.03 foci/cell for gMUC4-TwoMM, again demonstrating SNV-sensitivity and suggesting proteinase treatment is dispensable (Extended Data Fig. 6c). We also performed sgGOLDFISH against the *LMNA* gene using doubly mismatched (gLMNA-MUT) or singly mismatched (gLMNA-WT) guide RNA (Extended Data Fig. 7 and Fig. 8a). We observed 1.17 foci/cell for gLMNA-WT and 0.09 foci/cell for gLMNA-MUT (Fig. 1c). These data show that a single-nucleotide difference can produce about an order of magnitude difference in sgGOLDFISH detection efficiency. The full width at half maximum of Gaussian fit to the imaged *LMNA* foci is 617 ± 92 nm (mean ± SD, Extended Data Fig. 8b), indicating sgGOLDFISH provides well-defined spots suitable for subnuclear localization analysis.

We next applied sgGOLDFISH in patient-derived Hutchinson-Gilford progeria syndrome (HGPS) fibroblasts. HGPS cells contain one copy of normal *LMNA* gene (*LMNA*-WT), and one copy of mutated *LMNA* gene (*LMNA*-MUT) that carries a point mutation (c. 1824 C>T) (Fig. 2a), which causes expression of progerin, a truncated gene product, and alterations of nuclear shape^17^. The gLMNA-MUT guide RNA described above contains two mismatches against the wild-type *LMNA* sequence and one mismatch against the progeria mutant sequence (Extended Data Fig. 8c), and the gLMNA-WT has one mismatch against the wild-type and two mismatches against the mutant (Extended Data Fig. 8d). Therefore, sgGOLDFISH using gLMNA-MUT should preferentially label the mutant allele whereas the wild type allele is preferentially labeled when gLMNA-WT is used (Fig. 1a). To test this prediction, we created HGPS mutation-corrected fibroblasts by delivering adenine base editor ABE7.10max-VRQR (ABE) mRNA and corresponding sgRNA into the HGPS cells^18^ (Fig. 2b). This DNA-free approach efficiently corrected the HGPS mutation (> 94% efficiency) without the risk of unwanted DNA integration into the genome (Fig. 2c and Extended Data Fig. 9a). Consistently, the fraction of morphologically abnormal nuclei was significantly reduced after ABE treatment (Extended Data Fig. 9b and 9c).

**Fig. 2.**
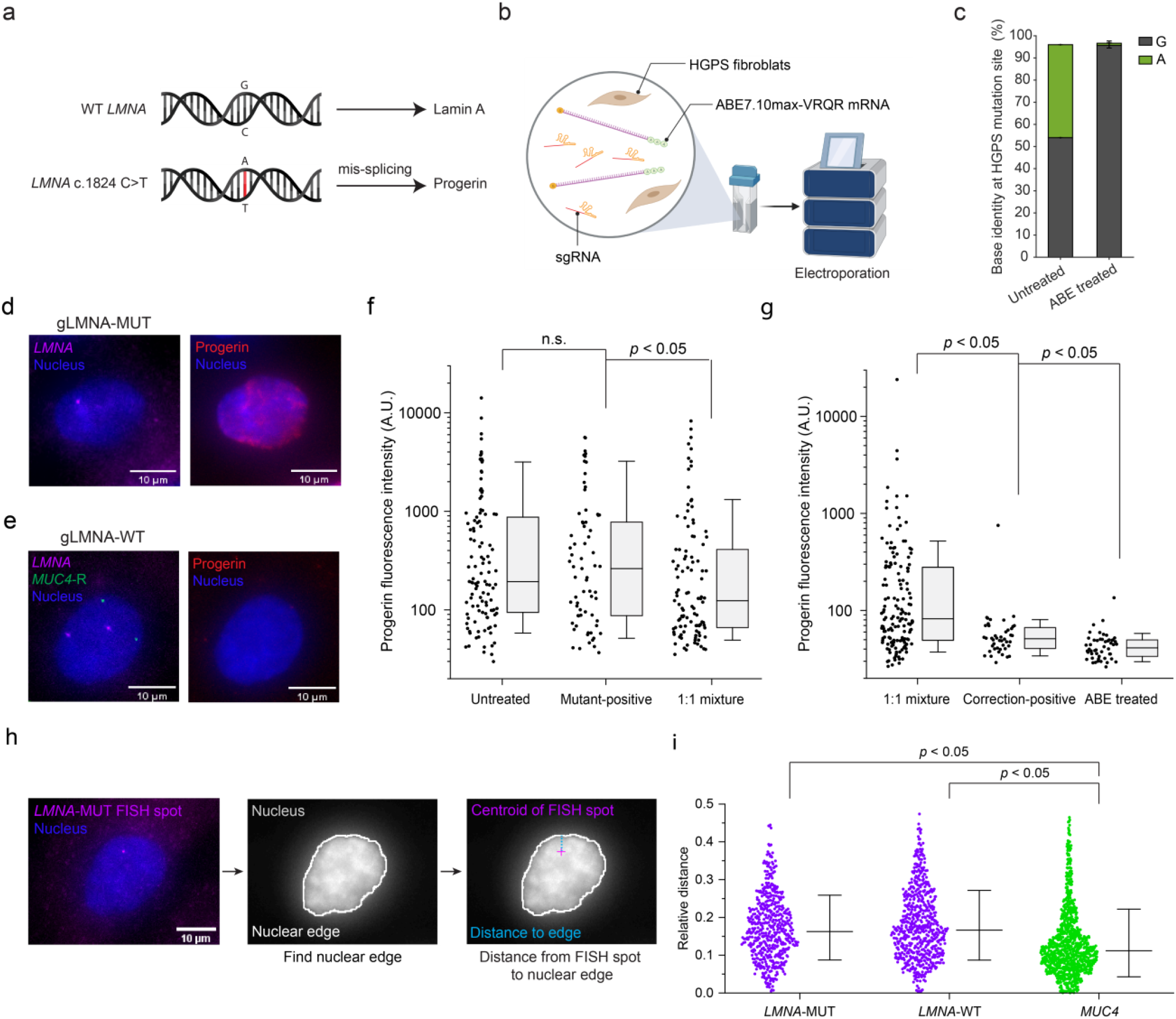
SNV detection in HGPS cells. **a**, Schematic of HGPS pathogenic point mutation. **b**, Schematic of ABE editing of HGPS fibroblasts. **c**, Base identity at the HGPS mutation site before and after ABE treatment. The error bar represents mean ± SD (n=3). **d** and **e**, sgGOLDFISH in parallel with progerin immunofluorescence using (**d**) gLMNA-MUT or (**e**) gLMNA-WT. **f** and **g**, Quantifications of progerin immunofluorescence intensity from different populations. Each dot represents a quantified cell (n=44 to 152 for each condition). Mann-Whitney *U* Test is used. n.s. represents *p* > 0.05. Box represents the range of 25^th^ to 75^th^ percentiles, and whisker represents the range of 10^th^ to 90^th^ percentiles. Median line is shown in the box. **h**, Schematic of measuring distance from a FISH spot to the nuclear edge. **i**, Quantifications of the relative distance (i.e., the distance from a FISH spot to the nuclear edge divided the square root of the nuclear area) of *LMNA*-WT, *LMNA*-MUT and *MUC4* alleles to the nuclear edge. Each dot represents a quantified FISH spot (n=538 to 994 for each condition). Student’s t test is used. Median line is shown. Whisker represents mean ± SD.

To test if sgGOLDFISH preferentially labels the progeria mutant allele with gLMNA-MUT, we made a cell mixture containing 50 % uncorrected HGPS cells and 50 % ABE-corrected HGPS cells (hereinafter called 1:1 mixture, Extended Data Fig. 9d). sgGOLDFISH against the *LMNA* gene using gLMNA-MUT was applied to the 1:1 mixture in parallel with progerin immunofluorescence, and a cell with at least one *LMNA* sgGOLDFISH spot was assigned as a “mutant-positive cell” (Fig. 2d and Extended Data Fig. 10a). The progerin immunofluorescence intensity averaged over the nucleus was comparable between the mutant-positive cells and untreated HGPS cells, but was significantly lower for cells randomly selected from the 1:1 mixture (Fig. 2f), consistent with reduced progerin expression after the HGPS mutation correction^18^. Therefore, sgGOLDFISH successfully identified uncorrected HGPS cells from a mixed population.

Next, we performed sgGOLDFISH in the 1:1 mixture again but using gLMNA-WT instead (Fig. 2e and Extended Data Fig. 10b). In parallel, we performed GOLDFISH against the *MUC4*-R region to simultaneously estimate the cell cycle stage (Fig. 2e, e.g., detection of two *MUC4*-R foci indicates G0/G1). We have previously shown the GOLDFISH detection efficiency of the *MUC4*-R region in fibroblasts is very high (around 90%)^13^. An ABE-corrected HGPS cell should have 2 to 4 copies of *LMNA*-WT alleles, depending on cell cycle. Therefore, cells with two *LMNA* foci and two *MUC4*-R foci, or four *LMNA* foci and four *MUC4*-R foci, were assigned as “correction-positive cells” (Fig. 2e). The progerin fluorescence was significantly lower for the correction-positive cells than for cells randomly selected from the 1:1 mixture (Fig. 2g). Taken together, these data suggest that sgGOLDFISH can preferentially label the *LMNA*-WT or the *LMNA*-MUT alleles even though these two alleles differ by only a single base pair, and that we can use sgGOLDFISH to identify base-edited cells or unedited cells from a heterogeneous population. We note that the progerin fluorescence is slightly higher for the correction-positive cells than for ABE-treated HGPS fibroblasts, likely because a small fraction of the *LMNA*-MUT alleles were labeled giving false positives (Fig. 2g).

Direct hybridization of probes allows for accurate localization of sequences of interest. To demonstrate sgGOLDFISH can facilitate sub-nuclear spatial analysis, we measured the distance of *LMNA*-WT, *LMNA*-MUT and *MUC4* foci to the nuclear edge in HGPS fibroblasts. *MUC4* foci appeared closer to the nuclear edge than *LMNA*-WT and *LMNA*-MUT foci (Fig. 2h and 2i), consistent with Lamin A/C-ChIP (chromatin immunoprecipitation) data from the same HGPS line which showed stronger Lamin A/C-ChIP signal for *MUC4* compared to *LMNA*, where Lamin A/C association correlates to a gene’s proximity to the nuclear membrane (Extended Data Fig. 10c)^19^.

Instead of labeling bulky amplification products in other methods^10, 11^ (e.g., the length of rolling circle amplification product varies from 160kb to 1Mb^20^), sgGOLDFISH labels 1kb to 1.5kb chromatin flanking the target SNV, which allows for accurate sub-nuclear localization of the target SNV. The much smaller labeling region of sgGOLDFISH may also explain that proteinase treatment is dispensable for sgGOLDFISH but the other methods require proteinase to digest cellular proteins for the amplification enzymes to function in the nucleus^10, 11^. Although the cleavage activity of Cas9 is dependent on local sequence, it can be fine-tuned by adding/removing rationally designed mismatches in the guide RNA or the 5’ extended guanine for new targets^15^. The limitations of our method include a requirement to test eCas9 RNP activity for new targets before sgGOLDFISH, slight nuclear shrinkage caused by fixation^13^, and ∼25% detection efficiency compared to existing SNV-sensitive FISH methods (10%-65%)^2, 3, 8-10^). Overall, given the single-nucleotide sensitivity, immunofluorescence compatibility, the ability to accurately locate SNVs, and relatively broad SNV targeting scope (see Supplementary Table 1 for comparison with other methods), sgGOLDFISH will be of value for researchers to study point mutation-related diseases or detect precise genome editing such as base editing.

## Methods

### Human Cell Lines

HEK293T cells were purchased from the American Type Culture Collection and cultured in DMEM with 4.5 g/L glucose, L-glutamine, and sodium pyruvate (Corning, 10-013-CV) supplemented with 10% heat inactivated fetal bovine serum (FBS, Corning 35-011-CV). Hutchinson-Gilford Progeria Syndrome (HGPS) fibroblasts were purchased from the Progeria Research Foundation and cultured in high glucose DMEM without L-glutamine (ThermoFisher, 11960-440) supplemented with 20% FBS (Corning, 35-011-CV), 1% Penicillin-Streptomycin (ThermoFisher, 15140-122) and 1% GlutaMAX (ThermoFisher, 35050-061). All cells were maintained at 37°C in 5% CO_2_ and imaging dishes were coated with 1 μg/cm^2^ fibronectin then air dried before plating.

### Expression and purification of Cas9 and Rep-X

Cas9 nickase and eCas9 nickase were prepared as described previously with modifications^21^. Cas9 nickase was expressed using the pMJ826 plasmid (addgene, 39316). Mutagenesis was carried out to introduce the H840A mutation into eSpCas9(1.1) variant using pJSC114 plasmid (addgene, 101215) and QuikChange Lightning Site-Directed Mutagenesis Kit (Agilent, 210518). eCas9 nickase was expressed using the mutagenesis-modified pJSC114 plasmid. The plasmids were transformed into NEB BL21(DE3) competent cells (New England Biosciences). Cultures were maintained in Terrific Broth (Invitrogen) supplemented with 0.4% glycerol at 37°C until induction at OD_600_ = 0.5 – 0.6 at which point the temperature was lowered to 18°C and cultures were induced with 0.5 mM IPTG (GoldenBio). Pellets were harvested after 16-18h and resuspended in lysis buffer (50 mM Tris-HCl, 500 mM NaCl, 5% glycerol, 1 tablets per 50 ml protease inhibitor (EDTA-free, Roche), 0.2 mM PMSF, 1 mM TCEP, 1 mg/ml lysozyme, pH 7.5) and sonicated at 30% amplitude with 2s on, 4s duty cycle for 2 min, 3 times. Lysate was spun down and supernatant was mixed with 2 ml Ni-NTA resin (Qiagen) per 50 ml sample and incubated for 1h at 4°C, then spun down and decanted. Resin was incubated with Wash Buffer (50 mM Tris-HCl, 500 mM NaCl, 10 mM imidazole, 5% glycerol, 1 mM TCEP, pH 7.5) at 4°C for 5 min repeated 4 times then added to gravity column. Colum was then incubated with Elution Buffer (50 mM Tris-HCl, 500 mM NaCl, 1 mM TCEP, 300 mM imidazole, 5% glycerol, pH 8 – 8.5) and fractions were analyzed via denaturing PAGE. Samples were then desalted and concentrated using a 40kD cut off filter into storage buffer (300 mM NaCl, 10 mM Tris-HCl, 0.1 mM EDTA, 50% glycerol, pH 7.5). Ni-NTA purification with desalting showed sufficient purity and activity for GOLD FISH applications and did not require further size selection chromatography.

Rep-X was prepared the same as previously described^13^. pET28a(+) with *rep* (C18L/C43S/C167V/C612A/S400C) was transformed into *E. coli* B21(DE3) (Sigma-Aldrich, CMC0014). A single colony was picked and grown in TB at 37°C overnight, followed by 30 °C overnight. When OD reached the range between 0.3 and 0.4, the cells were moved to an 18 °C incubator. When OD reaches 0.6 to 0.8, the cells were induced expression with 0.5 mM IPTG and continue growth overnight. The cells were harvested by centrifugation for 15 min at 5000 rpm and 4 °C. The pellet was resuspended in 40 ml of the lysis buffer (50 mM Tris-HCl pH 7.5, 5 mM Imidazole, 200 mM NaCl, 20% (w/v) sucrose, 15% (v/v) glycerol, 17.5 μg/ml PMSF, and 0.2 mg/ml Lysozyme) and sonicate to lyse the cells. The lysed cell mix was centrifuged at 14,000 rpm at 4 °C for 30-60 min and collect the supernatant. The supernatant was stir-mixed with pre-equilibrated Ni-NTA resin for 1.5 hours at 4 °C. Ni-NTA purification was performed by washing the protein-bound resin with buffer A (50 mM Tris-HCl pH 7.5, 5 mM Imidazole, 150 mM NaCl, 25% (v/v) glycerol), followed by buffer A1M (50 mM Tris-HCl pH 7.5, 5 mM Imidazole, 1 M NaCl, 25% (v/v) glycerol) to remove any DNA residue, and final washed the protein-bound resin with buffer A, then eluted the Rep variant with imidazole buffer (50 mM Tris-HCl pH 7.5, 205 mM Imidazole, 150 mM NaCl, 25% (v/v) glycerol). 20 µM eluted Rep variant was mixed with 100 µM BMOE crosslinker to self-crosslink into Rep-X. The reaction was stir-mixed at room temperature for 1 hour. The excess crosslinker and Imidazole was removed by an overnight dialysis and stored in Rep-X storage buffer (50% glycerol, 600 mM NaCl, 50 mM Tris-HCl pH 7.5) at -80 °C.

### Genome sequences

The GRCh38.p13 Primary Assembly was used in this study and downloaded from NCBI. The coordinates of target loci are listed below:

*MUC4*-R region (Chr3: 195788656-195778790)

*MUC4*-NR region (Chr3: 195807684-195808777)

*LMNA* region (Chr1: 156137082-156138607)

### sgGOLDFISH guide RNA and probe design

The SNV site should be within a protospacer of SpCas9. Because the previous study has demonstrated to image SNV at the PAM-proximal region^10^, here we focused on testing SNVs located at PAM-distal region. The 13^rd^ to 18^th^ positions from the PAM are ideal (Extended Data Fig. 2d). Because the eCas9 RNP can tolerate one PAM-distal mismatch, but two PAM-distal mismatches essentially inhibit cleavage under our conditions (Extended Data Fig. 2d, Extended Data Fig. 3c and Extended Data Fig. 7), an additional mismatch was intentionally introduced into the guide RNA (e.g., the U at the 8^th^ position from the 5’ end of crRNA in gMUC4-TwoMM and gMUC4-OneMM, Extended Data Fig. 2c). Oligo FISH probes for sgGOLDFISH were designed using Oligoarray^22^. The target DNA sequence (∼ 1.5 kb) immediately following the target protospacer was input into the Oligoarray 2.1 with the following constraints: Length: 18- to 24-nt; Tm: 70 °C to 90 °C; %GC: 30-70; Max. Tm for structure: 54 °C; Min. Tm to consider X-hybrid: 54 °C; and there was no consecutive repeat of 5 or more identical nucleotides. Probes that can non-specifically bind to human genome, human noncoding RNA and *E*.*coli* tRNA were removed. The probe sequences are listed in Supplementary Table 2.

### Preparation of DNA and RNA

The designed oligo FISH probes (without any labeling/modification) were purchased from IDT, and fluorescently labeled as previously described^23^. Briefly, to conjugate an amino-ddUTP at the 3’ end of each oligonucleotide, 66.7 µM DNA oligonucleotides, 200 µM Amino-11-ddUTP (Lumiprobe) and 0.4U/µl Terminal Deoxynucleotidyl Transferase (TdT, Thermo Scientific, EP0162) were mixed in 1X TdT Reaction buffer (Thermo Scientific) and incubated overnight at 37 °C. The reaction was cleaned up by ethanol precipitations and P4 beads (Bio-Rad, #1504124) spin columns. Next, the DNA oligonucleotides conjugated with amino-ddUTP were mixed with 100 µg of Cy3-NHS or Cy5-NHS (Lumiprobe or GE Healthcare) in 0.1 M sodium bicarbonate and incubated overnight at room temperature, and cleaned up by ethanol precipitations and P4 beads (Bio-Rad, #1504124) spin columns. Unlabeled oligonucleotides were removed by high-performance liquid chromatography (HPLC). The DNA substrates for *in vitro* cleavage assays were synthesized using Phusion^®^ Hot Start Flex 2X Master Mix (NEB, M0536S) and purified using GeneJET PCR Purification Kit (Thermo Scientific, K0701). The primers were purchase from IDT and sequences are listed in Supplementary Table 2. For the guide RNA against the *MUC4* gene, crRNA was synthesized *in vitro* using HiScribe™ T7 Quick High Yield RNA Synthesis Kit (NEB, E2050S) according to the manufacturer’s instructions, and purified by polyacrylamide gel electrophoresis. Alt-R^®^ CRISPR-Cas9 tracrRNA (IDT) was purchase from IDT. The guide RNA was annealed by mixing crRNA and tracrRNA at 1:1 ratio in Nuclease Free Duplex Buffer (IDT), and incubating at 95 °C for 30 seconds, then slowly cooling down to room temperature over 1 hour. For other guide RNAs used in this study, the guide RNA was synthesized using EnGen^®^ sgRNA Synthesis Kit, *S. pyogenes* (NEB, E3322V) according to the manufacturer’s instructions. The template DNA sequences are listed in Supplementary Table 2.

### *In vitro* cleavage assay

For Extended Data Fig. 2d, 2e and 7b, Cas9 RNP was assembled by mixing 200 nM eCas9 nickase with 400 nM guide RNA in the cleavage buffer (20 mM Hepes pH 7.5, 100 mM KCl, 7 mM MgCl_2_, 5% (v/v) glycerol and 0.1% (v/v) TWEEN-20, freshly added 1 mM DTT), and incubated for 10 min at room temperature. Then 4 nM DNA substrate was added, and incubated at 37 °C for 1 hour. Next, 80 unites/mL of proteinase K (NEB, P8107S) was added to the reaction, and incubated at 37 °C for 30 min. The reaction was directly loaded into the agarose gel for electrophoresis. For Extended Data Fig. 4b, 400 nM Cas9 nickase RNP cleaving the top strand (Extended Data Fig. 3a, Step 2) and 400 nM Cas9 nickase RNP cleaving the bottom strand (Extended Data Fig. 3a, Step 2) were assemble by mixing Cas9 nickase and corresponding guide RNA at 1:1 ratio and incubated for 10 min at room temperature. Then, 600 ng PCR-synthesized DNA substrate (Extended Data Fig. 2c) was added to the mixture and incubated for 1 hour at 37 °C. Next, 80 unites/mL of proteinase K (NEB, P8107S) was added to the reaction, and incubated at 37 °C for 30 min. The reaction was heated at 90 °C for 1 min to dissociate the two parts of the double-nicked DNA, followed by agarose gel electrophoresis.

### Cell fixation

The HEK293T or HGPS cells adhered to the glass surface of an imaging dish were fixed at -20 °C for 15 min in pre-chilled MAA solution (methanol and acetic acid mixed at 1:1 ratio), then washed three times (5 min each wash at room temperature unless indicated) with PBS. For SSB-ddPCR and the sgGOLDFISH in Fig. 1b, an additional protease treatment was performed: 0.1% pepsin in 0.1 M HCl was applied to the fixed HEK293T cells and incubated for 45 s at 37 °C, and the cells were washed with PBS once, and incubated in 70%, 90% and 100% EtOH at room temperature, each for 1 min. The cells were then washed three times with PBS.

### SSB-ddPCR

The SSB-ddPCR was performed similarly to DSB-ddPCR^16^ with modifications (Extended Data Fig. 3a). The fixed and pepsin treated HEK293T cells adhered to the glass surface of the imaging dish were incubated in the binding-blocking buffer (20 mM Hepes pH 7.5, 100 mM KCl, 7 mM MgCl_2_, 5% (v/v) glycerol and 0.1% (v/v) TWEEN-20, 1% (w/v) BSA, freshly added 1 mM DTT, freshly added 0.1 mg/ml *E*.*coli* tRNA) for 10 min at 37 °C. Next, 100 nM eCas9 nickase was mixed with 200 nM gMUC4-TwoMM or gMUC4-OneMM in the binding-blocking buffer, and incubated for 10 min at room temperature. The 100 nM eCas9 nickase RNP was then applied to the cells, and incubated for 45 min at 37 °C. After that, 2 mM ATP was supplied to the 100 nM eCas9 nickase RNP solution (i.e., the 100 nM eCas9 nickase RNP in the binding-blocking buffer supplied with 2 mM ATP), and incubated the cells in the solution for another 90 min at 37 °C, followed by PBS wash 3 times. To harvest genomic DNA from the cells, 60 mAU/mL proteinase K (Qiagen, 69504) diluted in PBS was applied to the cells and incubated for 30 min at 37 °C. The solution was collected from the imaging dish, and genomic DNA was extracted using the DNeasy Blood & Tissue Kits (Qiagen, 69504) by following manufacturer’s protocol. The extracted genomic DNA (less than 8 ng/μL) was further treated with 400 nM Cas9 nickase RNP using the corresponding guide RNA in 1X NEBuffer r3.1 (NEB, B7203S) for 1 hour at 37 °C. Next, 45 unit/mL proteinase K (NEB, P8107S) was added to the reaction and incubated for 30 min at 37 °C. The genomic DNA was purified using Genomic DNA Clean & Concentrator-10 (Zymo, D4011) and eluted in water. Finally, 20 to 50 ng the genomic DNA was mixed with 250 nM probes, 900 nM primers and 250 unit/mL Eael (NEB, R0508S) in 1X ddPCR Supermix for Probes (no dUTP) (Bio-Rad, 1863023). Droplets were created using Droplet Generation Oil for Probes, DG8 Gaskets, DG8 Cartridges, and QX200 Droplet Generator (Bio-Rad); Droplets were transferred to a 96-well PCR plate and heat-sealed using PX1 PCR Plate Sealer (Bio-Rad). PCR amplification was performed with the following conditions: 95 °C for 10 min, 40 cycles of (94 °C for 30 sec, 55 °C for 30 sec, 72 °C for 2 min), 98 °C for 10 min, 12 °C hold. Droplets were then individually scanned using the QX200 Droplet Digital PCR system (Bio-Rad). To generate the standard curve, gMUC4-OneMM and dCas9 (instead of eCas9 nickase) was applied to the fixed and pepsin treated HEK293T cells as described above, and the genomic DNA was harvested (Extended Data Fig. 5, Step 1). Half of the genomic DNA was treated with Cas9 nickase RNP as described above, which produces “ss-nicked genomic DNA”. Another half of the genomic DNA (less than 8 ng/μL) was treated with 0.2 unit/μL MseI (NEB, R0525S) for 1 hour at 37 °C, and MseI was deactivated by incubating the reaction 20 min at 65 °C. Following the deactivation, the Msel-treated genomic DNA was purified using Genomic DNA Clean & Concentrator-10 (Zymo, D4011) and eluted in water, which produces “ds-cut genomic DNA”. The “ss-nicked genomic DNA” and “ds-cut genomic DNA” were then mixed at different ratios as shown in standard curve in Extended Data Fig. 3c, and performed ddPCR using the same settings describe above.

### sgGOLDFISH

The cells adhered to the glass surface of an imaging dish were incubated in the binding-blocking buffer (20 mM Hepes pH 7.5, 100 mM KCl, 7 mM MgCl_2_, 5% (v/v) glycerol and 0.1% (v/v) TWEEN-20, 1% (w/v) BSA, freshly added 1 mM DTT, freshly added 0.1 mg/ml *E*.*coli* tRNA) for 10 min at 37 °C. Next, 100 nM eCas9 nickase was mixed with 200 nM guide RNA in the binding-blocking buffer, and incubated for 10 min at room temperature. For concurrent *MUC4*-R GOLDFISH (Fig. 1b), additional 20 nM eCas9 nickase and 40 nM gMUC4-R were also assembled in the binding-blocking buffer. After that, 2 mM ATP and 300 μM Rep-X was supplied to the 100 nM eCas9 nickase RNP solution (i.e., the 100 nM eCas9 nickase RNP in the binding-blocking buffer supplied with 2 mM ATP and 300 μM Rep-X), and incubated the cells in the solution for another 90 min at 37 °C, followed by PBS wash 3 times. Next, RNase Cocktail™ Enzyme Mix (Invitrogen, AM2286) was diluted 100 times in PBS and incubated with the cells for 1 hour at 37 °C. The cells were washed three times with PBS. The cells were then incubated in freshly made hybridization buffer (20% (v/v) formamide, 2X saline-sodium citrate (SSC), 0.1 mg/mL *E*.*coli* tRNA, 10% (w/v) dextran sulfate, 2 mg/mL BSA) for 10 min at room temperature. Fluorescently labeled oligo FISH probes (1 nM for *MUC4*-R, 2.5 nM per oligo FISH probe for *MUC4*-NR and *LMNA*, i.e., 57.5 nM and 90 nM final probe concentration for *MUC4*-NR and *LMNA*) in the hybridization buffer were applied to the cells and incubated for 1 hour at room temperature (repetitive targets) or 37 °C (non-repetitive targets). The cells were washed twice (10 min each wash) with wash buffer (25% formamide, 2X SSC) at 37 °C and once with PBS at room temperature for 5 min. One drop of Hoechst 33342 Ready Flow™ Reagent (Invitrogen, R37165) was mixed with 2 ml of PBS and incubated with the cells for 2 min at room temperature. Finally, imaging buffer (2X SSC and saturated Trolox (> 5 mM), 0.8% (w/v) dextrose) supplied with GLOXY (1 mg/mL glucose oxidase, 0.04 mg/mL catalase) was added to the cells for imaging.

### Immunofluorescence

For progerin immunofluorescence, the cells after sgGOLDFISH were incubated in IF buffer (1X Blocker™ BSA in PBS (Thermo Scientific, 37525) supplied with 0.1% Tween-20) at room temperature for 20 min. Progerin Monoclonal Antibody (13A4) (Thermo Scientific, 39966) was diluted 500 times in the IF buffer, and applied to the cells for overnight incubation at 4 °C. The cells were washed three times with PBS, and incubated with 500 times diluted Alexa750-labeled secondary antibody (Invitrogen, A-21037) in the IF buffer for 30 min at room temperature. Finally, the cells were wash 3 times with PBS and imaged in the imaging buffer. For Lamin A/C immunofluorescence, the cells after fixation were incubated in IF buffer (1X Blocker™ BSA in PBS (Thermo Scientific, 37525) supplied with 0.1% Tween-20) at room temperature for 20 min. Anti-Lamin A + Lamin C antibody [4C11] (Abcam, ab238303) was diluted 500 times in the IF buffer, and incubated with the cells for 1 hour at room temperature. The cells were washed three times with PBS, and incubated with 500 times diluted Alexa750-labeled secondary antibody (Invitrogen, A-21037) in the IF buffer for 30 min at room temperature. Finally, the cells were wash 3 times with PBS and imaged in the imaging buffer.

### Microscopy

sgGOLDFISH imaging was performed using Nikon Eclipse Ti microscope equipped with Nikon perfect focus system, Xenon arc lamp. The system was driven by Elements software. Nikon 60X/1.49 NA objective (CFI Apo TIRF) was used. Emission was collected using a custom laser-blocking notch filter (ZET488/543/638/750M) from Chroma. Images were recorded using an electron-multiplying charge-coupled device (Andor iXon 888). Images were recorded as z-stacks (20 to 30 steps), with 300 nm to 500 nm step size.

### DNA-free base editing in HGPS fibroblasts

The guide RNA for base editor to correct the HGPS mutations (LMNA-VRQRABE-sgRNA) and DNA primers for preparing VRQR-ABE7.10max mRNA were purchased from IDT (see Supplementary Table 2 for the RNA and DNA sequences). To prepare DNA template for VRQR-ABE7.10max mRNA, pUC19 (NEB, N3041S) was linearized using EcoRI-HF (NEB, R3101S) and HindIII-HF (NEB, R3104S). VRQR-AEB fragment was PCR-synthesized using the Plasmid #154429 (addgene) and VRQR-AEB-primer-F and VRQR-AEB-primer-R, and gel purification was carried out to remove non-specific products. Mutagenesis was performed using pcDNA3.3-eGFP (addgene, Plasmid #26822) and T7-Mutagenesis-primer-F and T7-Mutagenesis-primer-R to replace the “G” following the T7 promoter sequence with an “A” by the QuikChange Lightning Site-Directed Mutagenesis Kit (Agilent, 210518). T7-5’UTR fragment was PCR-synthesized using the mutagenesis-modified pcDNA3.3-eGFP and T7-5’UTR-primer-F and T7-5’UTR-primer-R. The 3’UTR fragment was PCR-synthesized using the mutagenesis-modified pcDNA3.3-eGFP and 3’UTR-primer-F and 3’UTR-primer-R. Next, the linearized pUC19, VRQR-AEB fragment, T7-5’UTR fragment, 3’UTR fragment was assembled into a plasmid (VRQRABE-mRNA plasmid) using NEBuilder HiFi DNA Assembly Master Mix (NEB, E2621S) according to manufacturer’s protocol. The linear VRQRABE-mRNA DNA template was PCR-synthesized using VRQRABE-mRNA plasmid, VRQRABE-mRNA-linearTemplate-F and VRQRABE-mRNA-linearTemplate-R. All PCR reactions were performed using Q5^®^ Hot Start High-Fidelity 2X Master Mix (NEB, M0494S). The *in vitro* transcription of VRQR-ABE7.10max mRNA reaction contains 50 ng/μL linearized VRQRABE-mRNA DNA template, ATP/CTP/GTP (5 mM each), 5 mM N1-methylpseudouridine (TriLink, N-1081-1), 4 mM CleanCap AG (TriLink, N-7113), 1 unit/μL Murine RNase Inhibitor (NEB, M0314S), 0.002 units/μL Yeast Inorganic Pyrophosphatase (NEB, M2403S) and 8 units/μL T7 RNA Polymerase (NEB, M0251S) in the transcription buffer (40 mM Tris-HCl pH 8, 20 mM spermidine, 0.02% (v/v) Triton X-100, 165 mM magnesium acetate, freshly added 10 mM DTT). The *in vitro* transcription reaction was incubated at 37 °C for 2 hours, and treated with DNase I by supplying with 1X DNase buffer (NEB, B0303S) and 0.3 units/μL DNase I (M0303S) and incubating at 37°C for 20 min. The reaction was purified using Megaclear™ Transcription Clean-Up Kit (Invitrogen, AM1908), and dephosphorylated using 0.25 units/μL Antarctic Phosphatase (NEB, M0289S) according to manufacturer’s protocol. The VRQR-ABE7.10max mRNA was purified again using the Megaclear™ Transcription Clean-Up Kit. All electroporation experiments were carried out using the Lonza 4D-Nucleofector System. For mRNA editing in HGPS cells, 5 μg of LMNA-VRQRABE-sgRNA was mixed in a total 25 μL volume (SE kit, Lonza) then resuspended with 200k HGPS cells and electroporated using the with CM-120 setting. Cells were maintained at 37°C in 5% CO_2_ for 3 days before collecting genomic DNA using DNeasy Blood & Tissue Kits (Qiagen, 69504) and sequencing.

### Data analysis for GOLD FISH

Images were processed using Fiji/ImageJ. Z-stack images were projected to a single plane using the ‘Max Intensity’ Z-Projection function. The contrasts of images were linearly adjusted by changing the minimum and maximum values using the ‘brightness/contrast’ function in ImageJ for optimal visualization purpose only. FISH-quant was used to find foci in each cell and fitted with three-dimensional (3D) Gaussian function^24^. The nuclear edge, nuclear area and the distance from a FISH focus to the nuclear edge were analyzed using custom-written MATLAB scripts.

## Data availability

The data that support the findings of this study will be available at Mendeley data.

## Code availability

Any custom codes used in the study will be available at Github.

## Acknowledgements

We thank the Doudna laboratory (University of California-Berkeley) for generously providing dCas9 stocks. We thank the Regot lab (Johns Hopkins School of Medicine) for generously providing HEK293ft cell line. We thank the support of Worod Allak (Johns Hopkins School of Public Health) on ddPCR experiments. The project was supported by grants from the National Institutes of Health (GM122569 to T.H. and GM097330 to S.B) and the National Science Foundation [PHY-1430124 to T.H.]. T.H. is an investigator with the Howard Hughes Medical Institute.

## Author contributions

Y.W. and T.H. designed the experiments. Y.W. performed sgGOLDFISH and ddPCR experiments. R.Z. assisted ddPCR experiments. Y.W. synthesized ABE mRNA. T.C. cultured cells and performed electroporation. H.W. synthesized and purified crRNA. M.G. and T.C. expressed and purified proteins. Y.W. and T.C. performed data analysis. Y.W., T.H. and T.C. discussed the data. Y.W., T.H., T.C. and S.B. wrote the manuscript.

## Competing interests

The authors declare no competing interests.

**Extended Data Fig. 1.**
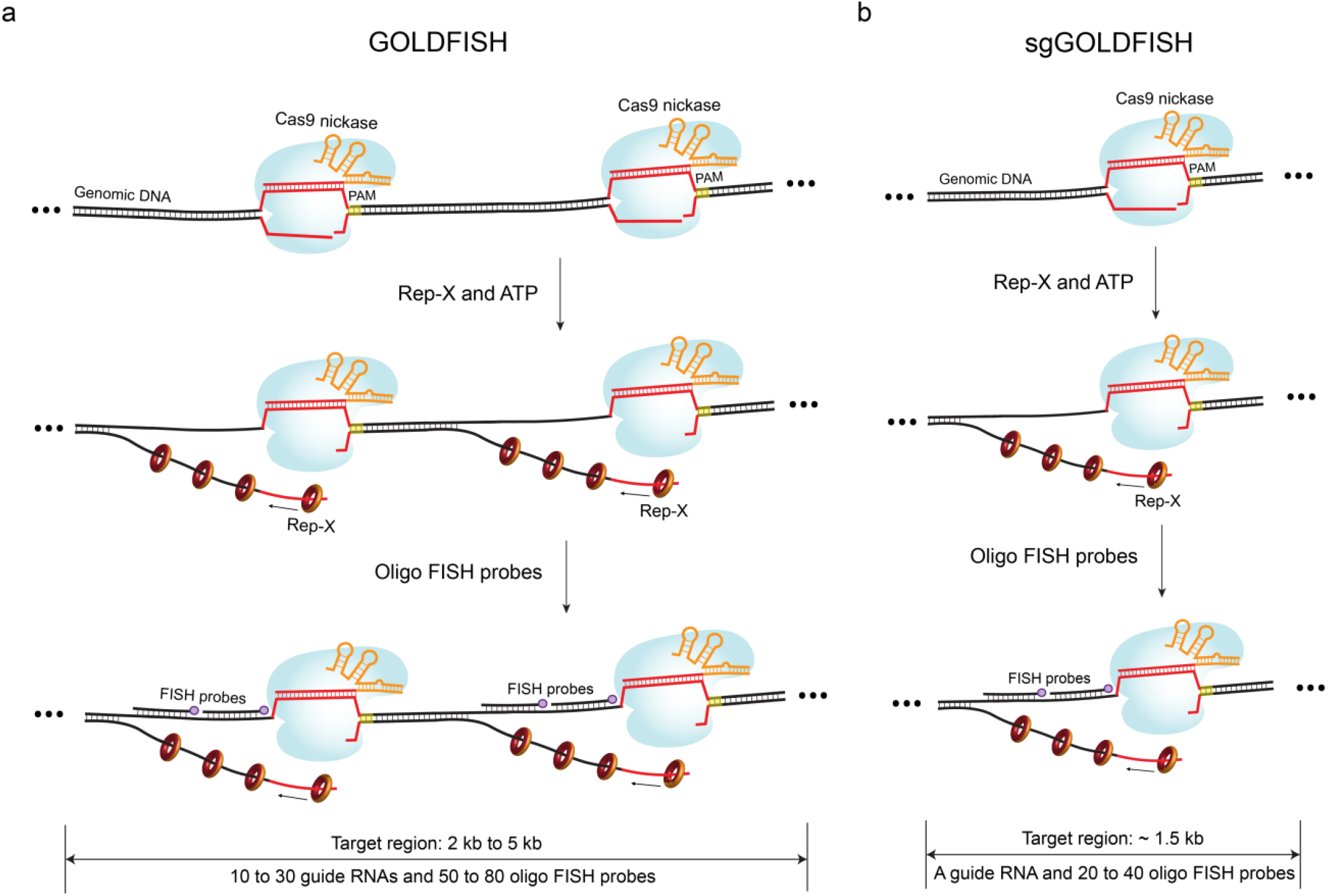
GOLDFISH and sgGOLDFISH. **a**, Schematic of GOLDFISH. Cas9 nickase RNP is applied to fixed and permeabilized cells to cleave the genomic DNA. Then Rep-X along with ATP is added to unwind the genomic DNA from the Cas9 cleavage sites. Finally, fluorescently labeled FISH probes are added to hybrid to sequences of interest. Multiple different guide RNA species and FISH probes are used in the GOLDFISH. The target region (i.e., guide RNA and probe binding sites) spans typically 2 kb to 5 kb. **b**, Schematic of sgGOLDFISH. The experimental procedure is the same as GOLDFISH, but only 1 guide RNA species is used in the sgGOLDFISH. The target region spans typically 1 kb to 1.5 kb.

**Extended Data Fig. 2.**
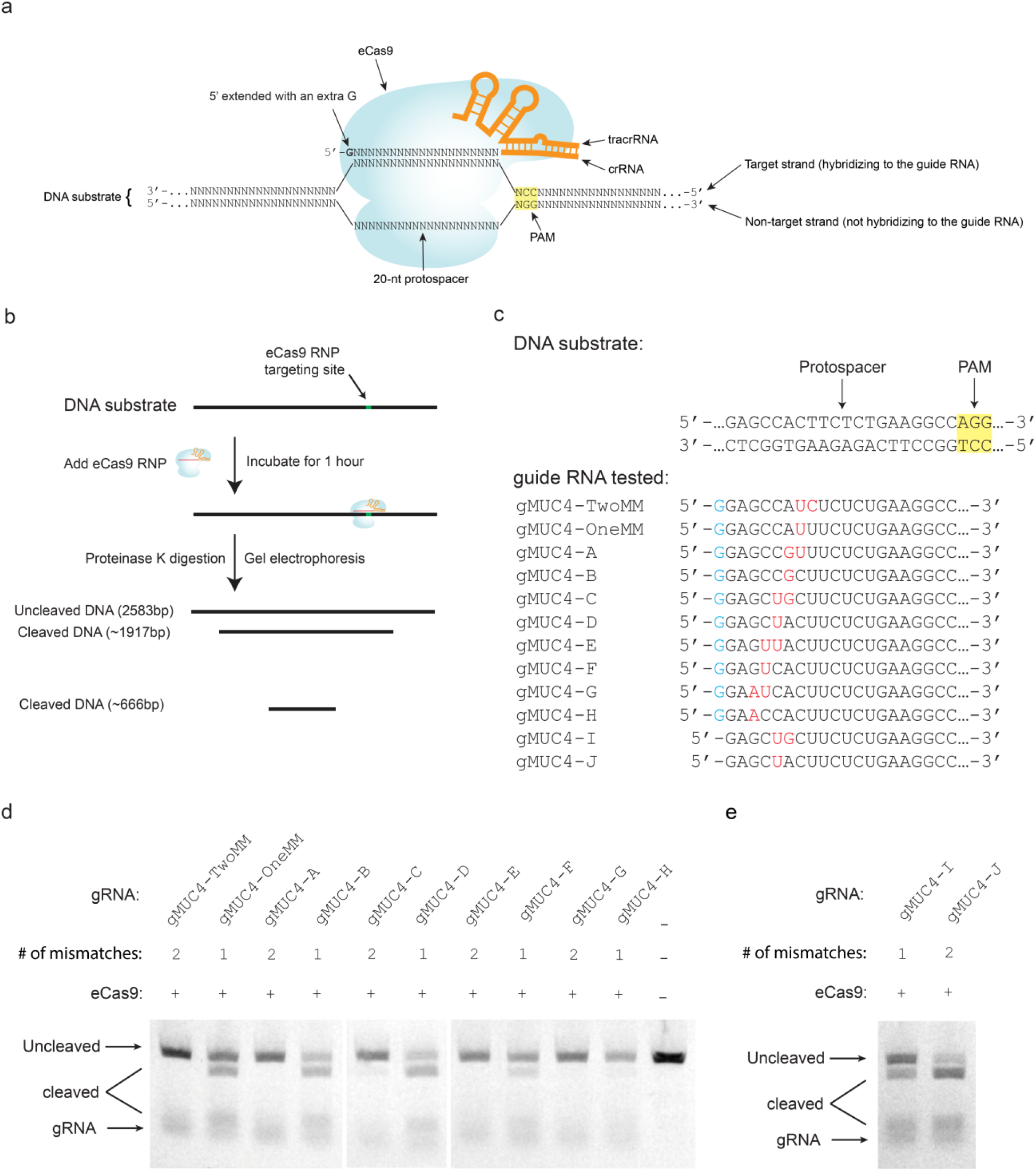
eCas9 RNP and the *in vitro* cleavage assay. **a**, Schematic of eCas9 RNP. Compared to canonical guide RNA, the 5’ extended guide RNA used in this study has an extra guanine (bolded in the figure) at the 5’ of crRNA. **b**, Schematic of *in vitro* cleavage assay. eCas9 RNP was mixed with DNA substrate and incubated for 1 hour at 37 °C. Then proteinase K was added to digest bound and free eCas9. Finally, the reaction was loaded into an agarose gel for electrophoresis. **c**, Sequences of DNA substrate and guide RNA tested (only 20 or 21 nucleotides from the 5’ end of the crRNA are shown) in the *in vitro* cleavage assay. The DNA substrate was PCR-synthesized using human genomic DNA and primers against a non-repetitive region of the *MUC4* gene. A group of guide RNAs with 1 or 2 mismatches against the target protospacer were used in the cleavage assay. The blue “G” represents the 5’ extended guanine of the crRNA. The red colored nucleotides represent mismatches against the DNA substrate. **d**, Gel image of the *in vitro* cleavage assay using guide RNA with 5’ extended guanine. **e**, Gel image of the *in vitro* cleavage assay using canonical guide RNA (i.e., without the 5’ extended guanine). Significant cleavage activity was observed with the canonical guide RNA even if there are two mismatches between the guide RNA and DNA substrate.

**Extended Data Fig. 3.**
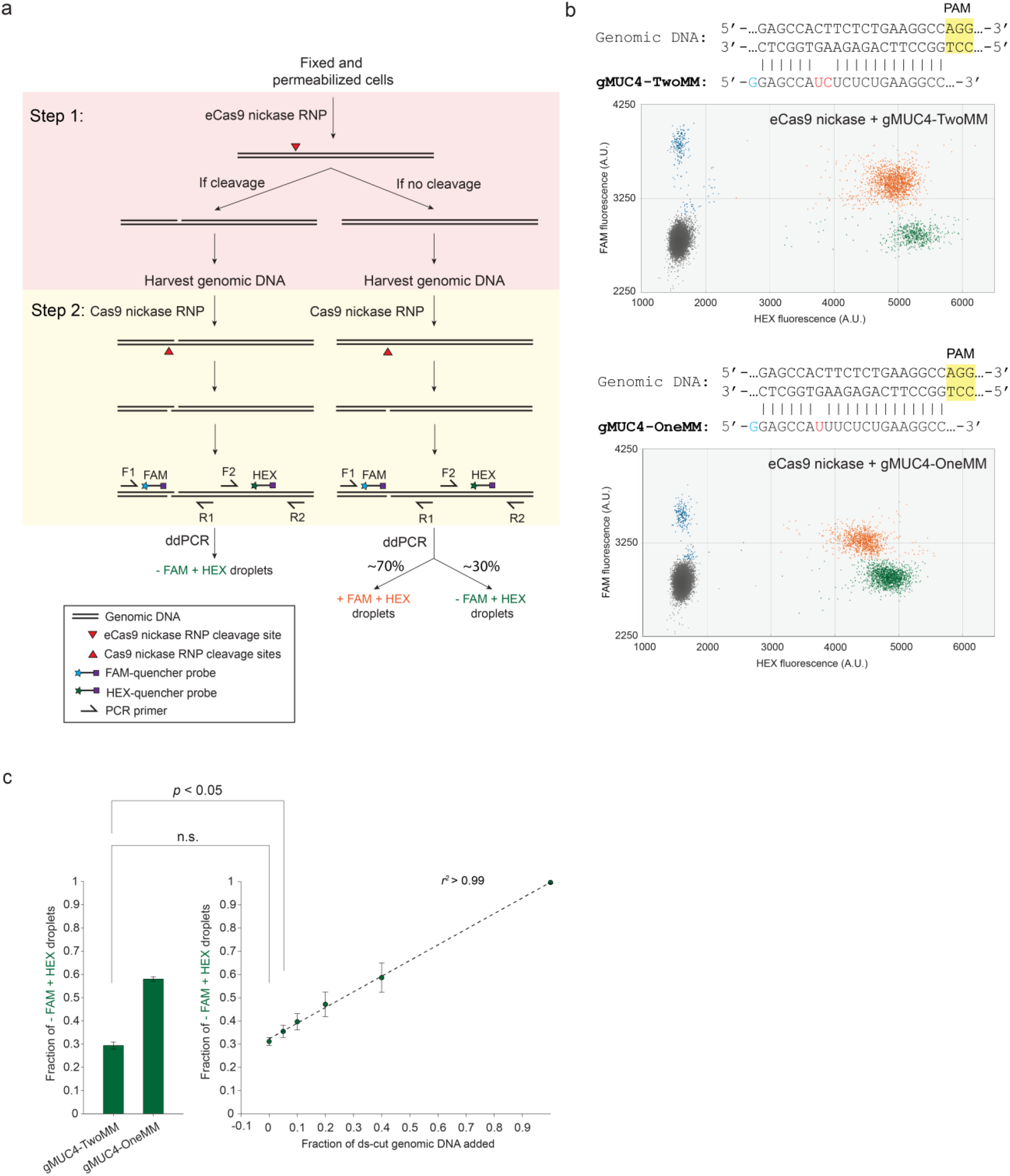
SSB-ddPCR. **a**, Schematic of SSB-ddPCR. See Supplementary Note for detailed description. **b**, Representative SSB-ddPCR scattering plot results using eCas9 nickase and gMUC4-TwoMM or gMUC4-OneMM. The 5’ sequences of gMUC4-TwoMM and gMUC4-OneMM as well as their target genomic DNA sequence are shown above each scattering plot. **c**, Left, bar plot of fraction of – FAM + HEX droplets from the SSB-ddPCR using gMUC4-TwoMM or gMUC4-OneMM. Right, standard curve of the ddPCR assay. Student’s t test is used. n.s. represents *p* > 0.05. Error bar represents mean ± standard deviation from at least 3 replicates.

**Extended Data Fig. 4.**
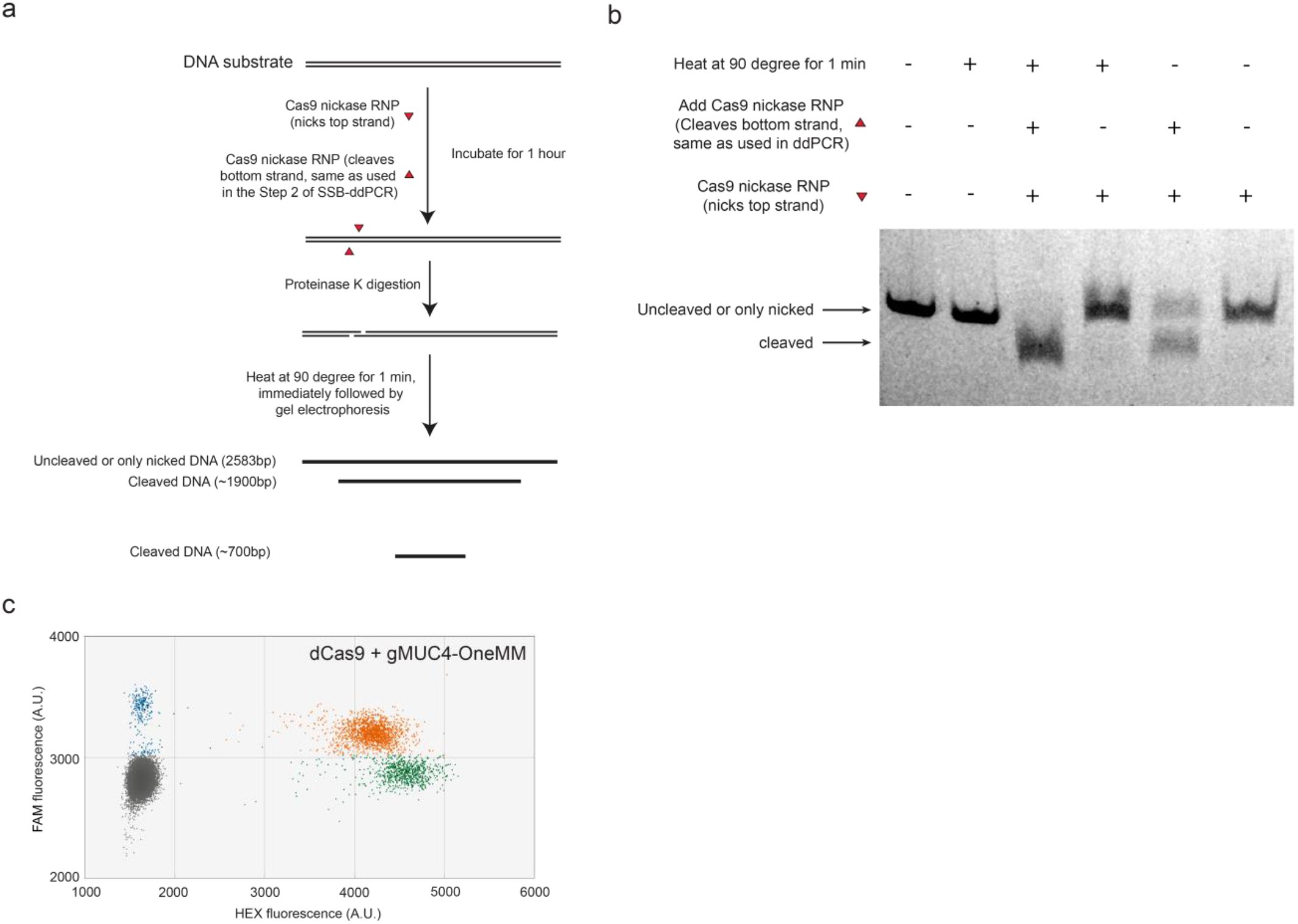
Control experiments for SSB-ddPCR. **a**, *In vitro* cleavage assay to measure the efficiency of DNA cleavage by Cas9 nickase RNP in the step 2 in Extended Data Fig. 3a. In this assay, less than 8 ng/μL PCR-synthesized DNA substrate (Extended Data Fig. 2c) was mixed with 400 nM Cas9 nickase RNP cleaving the top strand and 400 nM Cas9 nickase RNP cleaving the bottom strand, and incubated for 1 hour at 37 °C. The 400 nM Cas9 RNP cleaving the bottom strand was also used in the Step 2 in Extended Data Fig. 3a. After proteinase K treatment, the reaction was heated at 90 °C for 1 min to dissociate the two parts of the double-nicked DNA, followed by agarose gel electrophoresis. **b**, Gel image of the *in vitro* cleavage assay. Only the 3^rd^ lane shows close to 100% cleavage efficiency indicates the 400 nM Cas9 RNP cleaved almost all DNA molecules. **c**, Representative SSB-ddPCR result using dCas9 and gMUC4-OneMM.

**Extended Data Fig. 5.**
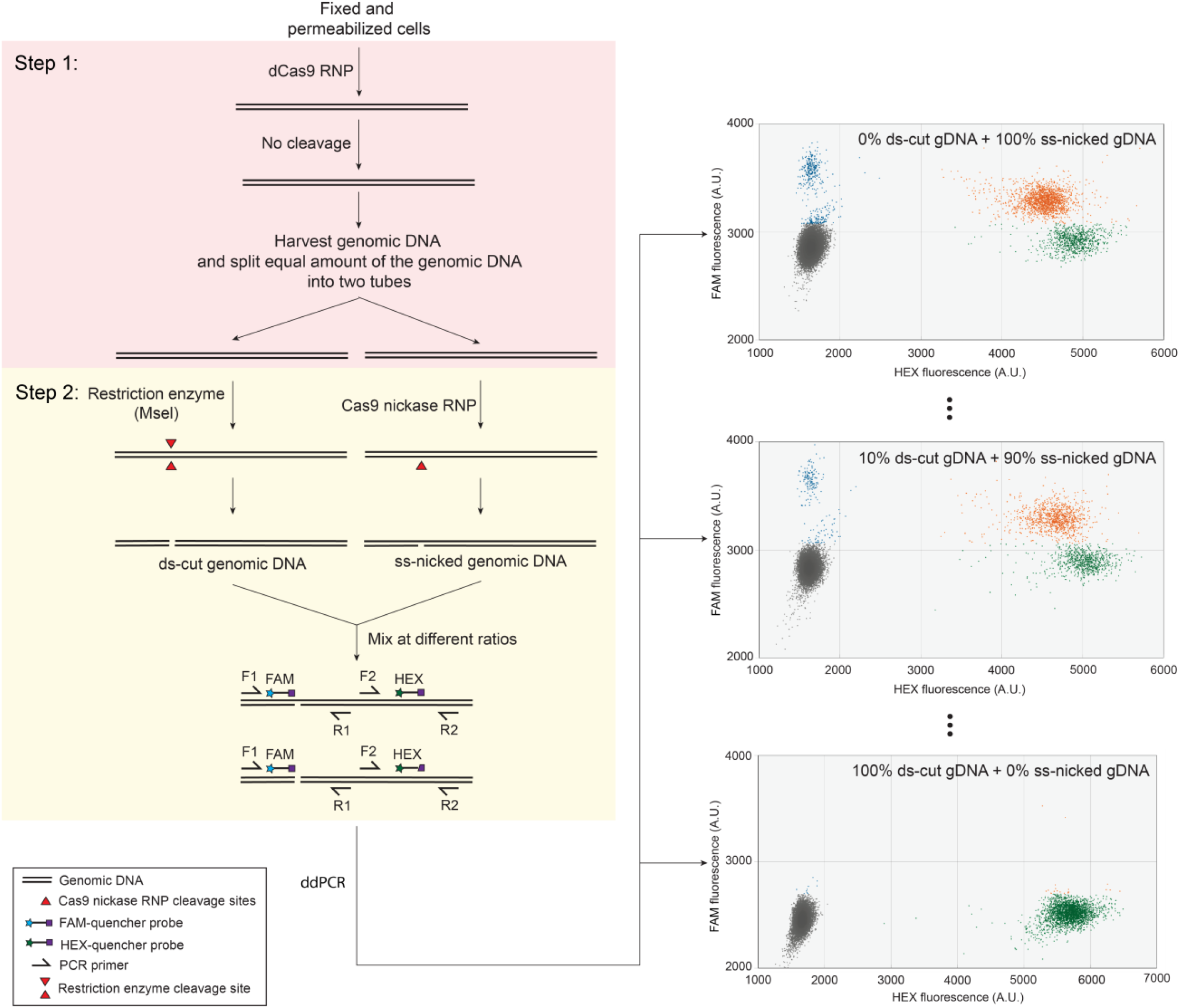
Generating standard curve of SSB-ddPCR. Schematic of the generating standard curve in Extended Data Fig. 3c. See Supplementary Note for detailed description.

**Extended Data Fig. 6.**
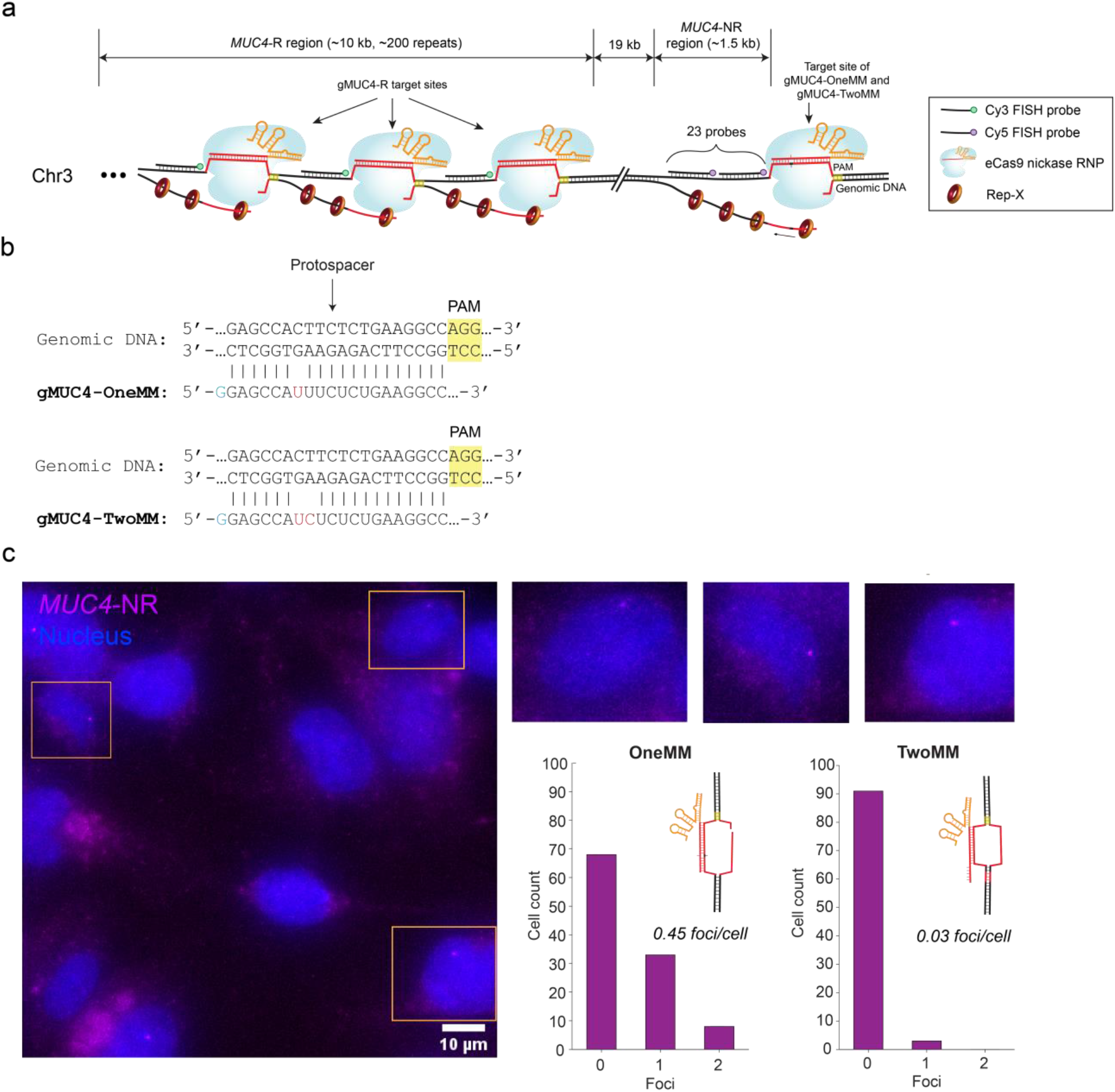
Schematic of sgGOLDFISH against the *MUC4*-NR region and GOLDFISH against the *MUC4*-R region. **a**, Only one guide RNA (gMUC4-OneMM or gMUC4-TwoMM) and 23 FISH probes were used to target the *MUC4*-NR region. The figure shows the case that gMUC4-OneMM was used (there is one mismatch between guide RNA and target DNA). In contrast, although only one guide RNA (gMUC4-R) and one FISH probe species were used to target *MUC4*-R region, the *MUC4*-R region contains ∼200 repeats, therefore multiple binding sites for eCas9 nickase RNP complexed with gMUC4-R and the FISH probe against *MUC4*-R region. **b**, Sequences of *MUC4*-NR target protospacer and (top) gMUC4-OneMM or (bottom) gMUC4-TwoMM (only 21 nucleotides from the 5’ end of the crRNA are shown). The blue-colored G represents the extended guanine at the 5’ of the guide RNA. Red-colored nucleotides represent mismatches. **c**, A representative sgGOLDFISH image using gMUC4-OneMM without proteinase treatment in HEK293T cells (single cells outlined in orange are magnified on the upper– right corner), and histograms of sgGOLDFISH foci using gMUC4-OneMM (n=109) or gMUC4-TwoMM (n=94).

**Extended Data Fig. 7.**
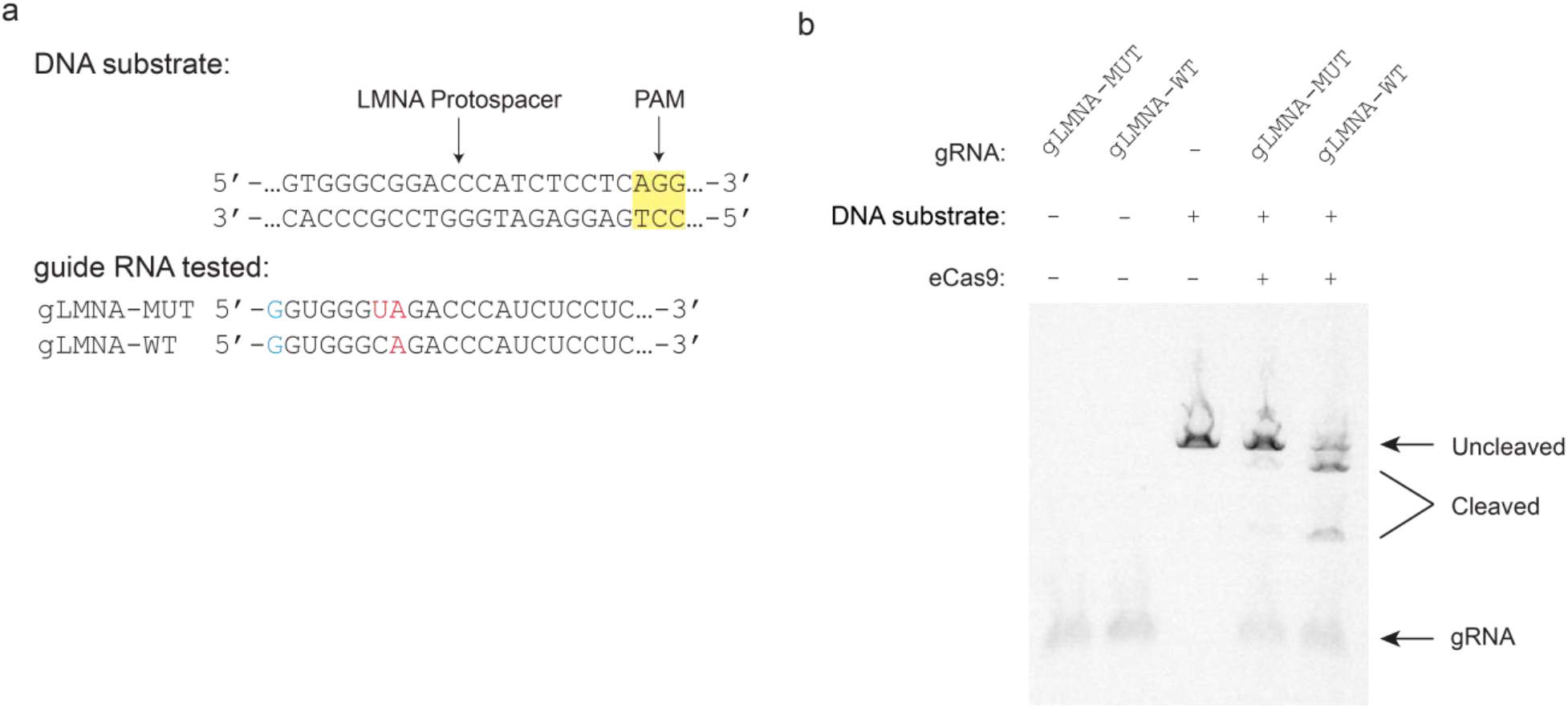
*In vitro* cleavage assay to measure cleavage activity of eCas9 in complex with gLMNA-MUT and gLMNA-WT against the *LMNA* gene. **a**, The DNA substrate was PCR-synthesized using human genomic DNA and primers against a non-repetitive region of the *LMNA* gene. The sequences of 5’ end of gLMNA-MUT and gLMNA-WT are shown. The blue “G” represents the 5’ extended guanine of the crRNA. The red colored nucleotides represent mismatches against the DNA substrate. **b**, Gel image of the *in vitro* cleavage assay using the guide RNAs and the DNA substrate in Extended Data Fig. 7a.

**Extended Data Fig. 8.**
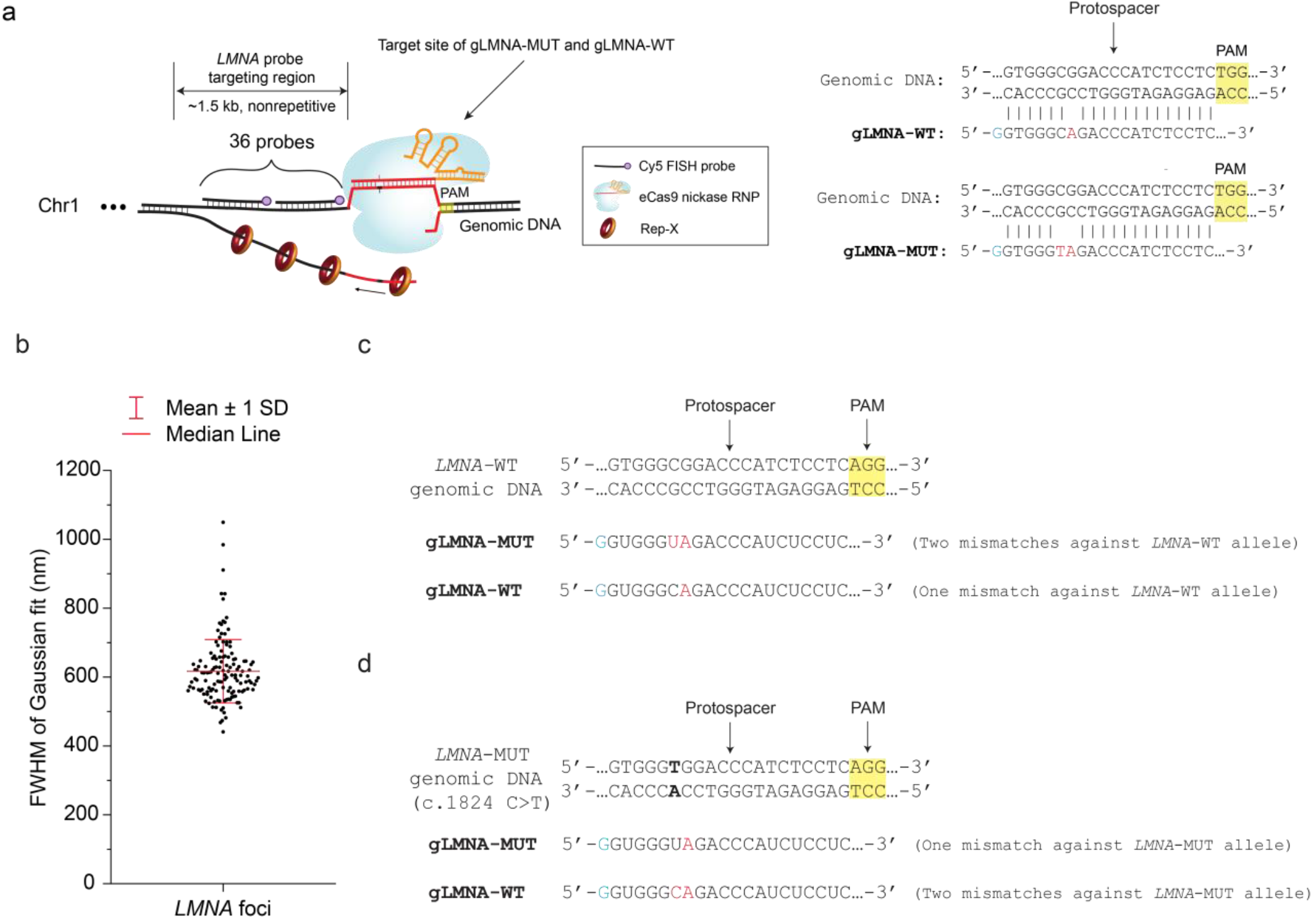
Schematics of sgGOLDFISH against *LMNA*. **a**, Left, schematic of sgGOLDFISH against *LMNA* using gLMNA-MUT or gLMNA-WT. The figure shows the scenario that gLMNA-WT is used to target a wild-type *LMNA* allele (there is one mismatch between guide RNA and target protospacer). Right, Sequences of target LMNA protospacer in HEK293T cells, gLMNA-WT and gLMNA-MUT (which were tested in Extended Data Fig. 7). The blue-colored G represents the extended guanine at the 5’ of the guide RNA. Red-colored nucleotides represent mismatches. **b**, Statistics of the full width at half maximum (FWHM) of the *LMNA* foci from the sgGOLDFISH using gLMNA-WT in Fig. 1c. Each dot represents a quantified *LMNA* focus (n=137). Median line is shown. Whisker represents mean ± standard deviation (SD). **c**, Sequences of the target protospacer of the *LMNA*-WT allele in HGPS cells and gLMNA-MUT or gLMNA-WT. The blue “G” represents the 5’ extended guanine of the guide RNA. The red colored nucleotides represent mismatches against the protospacer. **d**, Sequences of the target protospacer of the *LMNA*-MUT allele in HGPS cells and gLMNA-MUT or gLMNA-WT. The blue “G” represents the 5’ extended guanine of the guide RNA. The red colored nucleotides represent mismatches against the protospacer. The bolded ‘A-T’ base pair indicates the *LMNA* c.1824 C>T mutation in HGPS fibroblasts.

**Extended Data Fig. 9.**
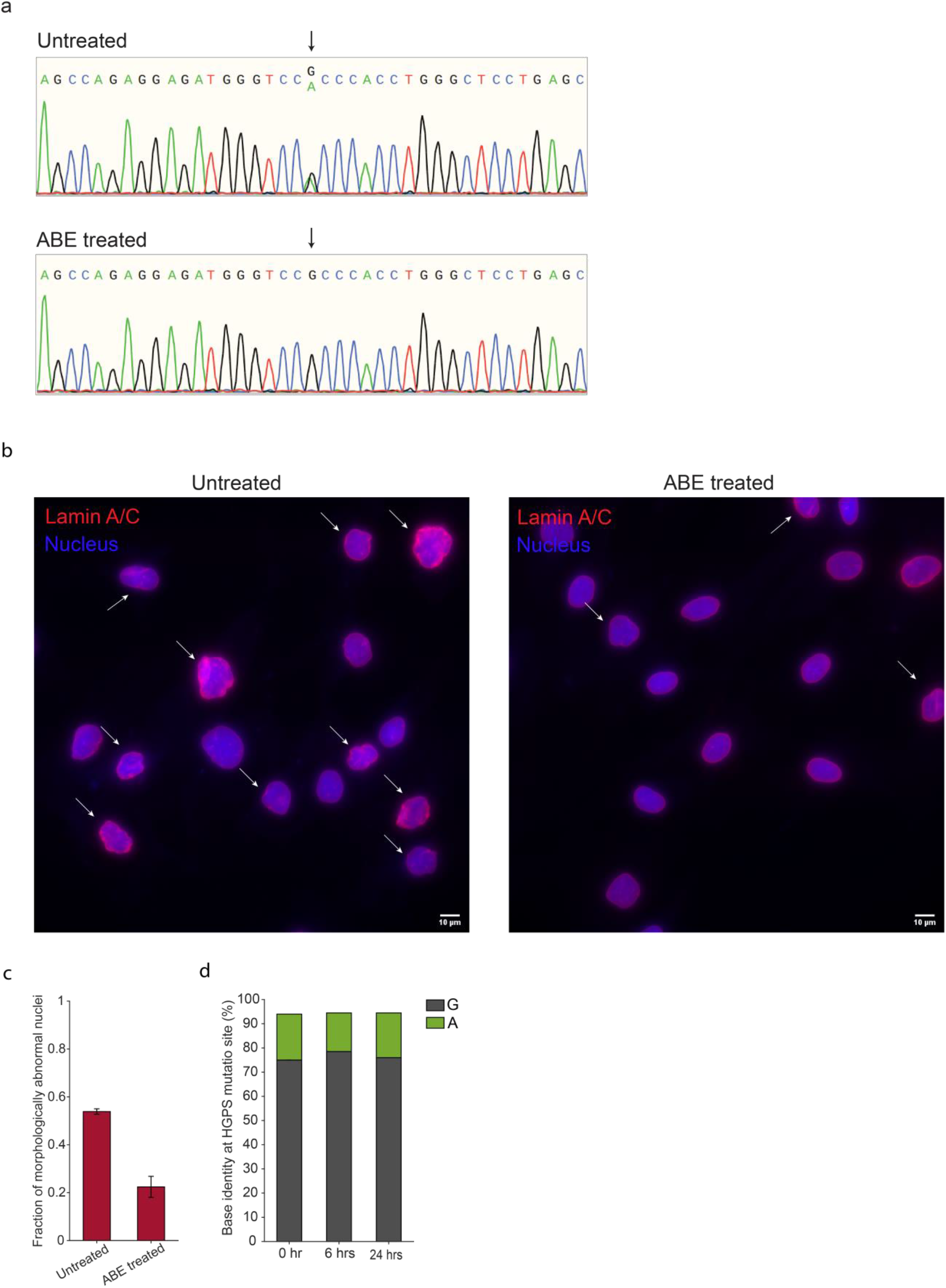
DNA-free base editing to correct the HGPS pathogenic point mutation. **a**, Representative Sanger sequencing results of untreated and ABE-treated HGPS fibroblasts. The black arrow indicates the HGPS pathogenic point mutation site. **b**, Representative images showing the morphology difference of Lamin A/C meshwork between untreated and ABE-treated HGPS fibroblasts. White arrows indicate morphologically abnormal nuclei. **c**, Quantification of fraction of morphologically abnormal nuclei in untreated and ABE-treated HGPS fibroblasts. Morphologically abnormal nuclei were identified by visual inspection. More than 150 cells were quantified for each condition. The dataset was quantified twice by two persons independently. The error bar indicates the difference between the two quantifications. Student’s t test is used. **d**, Quantifications of base identity at the HGPS pathogenic point mutation site at different time points after mixing untreated and ABE-treated HGPS fibroblasts at 1:1 ratio (i.e., 1:1 mixture). Base identity was measure by Sanger sequencing. The data indicate the 1:1 mixture contains roughly 50% uncorrected and 50% ABE-corrected HGPS fibroblast within 24 hours.

**Extended Data Fig. 10.**
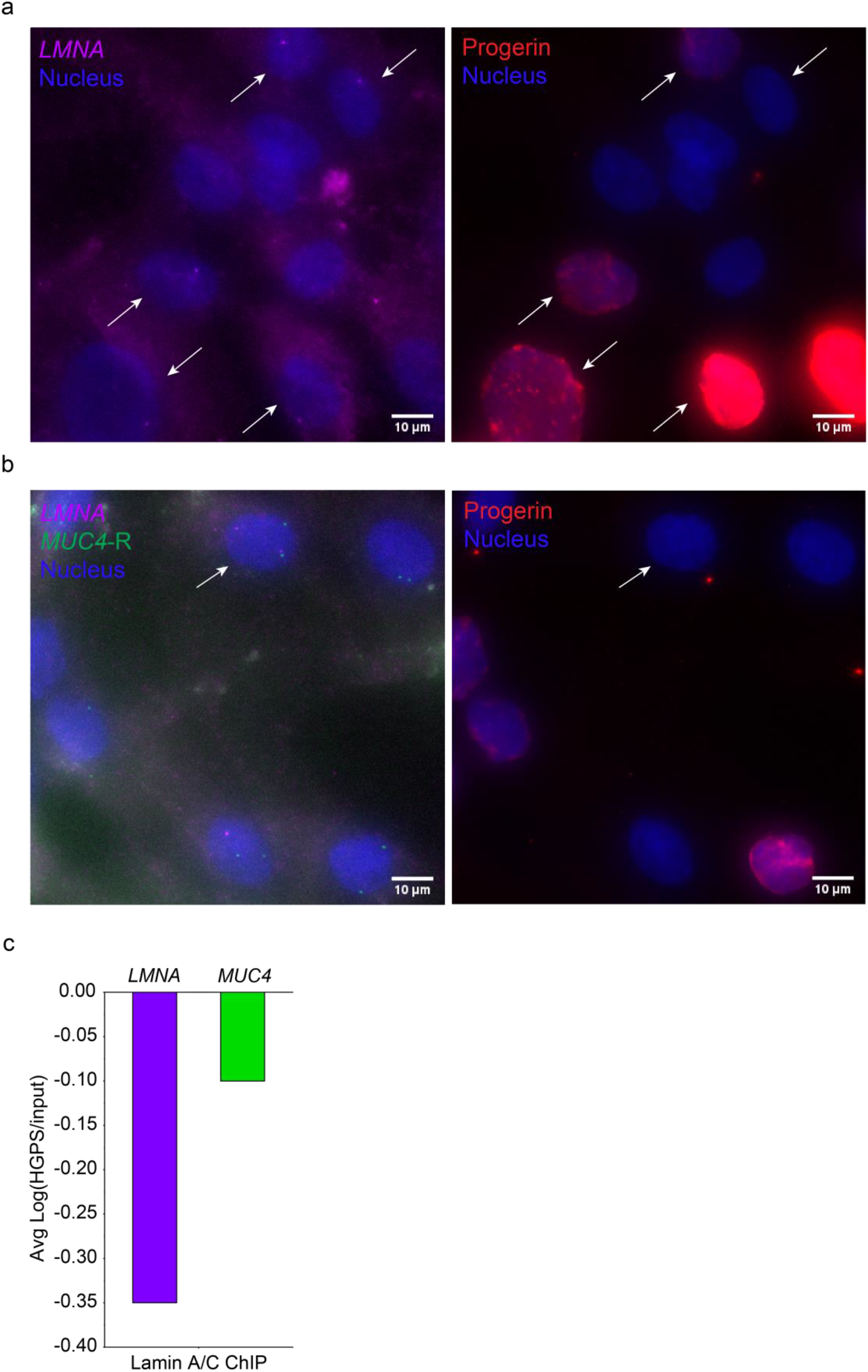
sgGOLDFISH signals can be used for spatial analysis. **a**, A representative image of sgGOLDFISH against *LMNA* using gLMNA-MUT and progerin immunofluorescence in 1:1 mixture. The “mutant-positive cells” are indicated by white arrows. **b**, A representative image of sgGOLDFISH against *LMNA* using gLMNA-WT and progerin immunofluorescence in 1:1 mixture. The “correction-positive cell” is indicated by a white arrow. **c**, Lamin A/C-ChIP data of the HGPS fibroblasts from a previous study (https://research.nhgri.nih.gov/manuscripts/Collins/HGPSepigenetics/). The enrichment (i.e., Avg Log(HGPS/Input)) of the *LMNA* and *MUC4* genes in Lamin A/C ChIP is shown. Smaller value (i.e., more negative) suggests less enriched in Lamin A/C ChIP. The data show that the *LMNA* gene is less enriched than the *MUC4* gene in Lamin A/C ChIP. This is consistent with our data in Fig. 2i showing that the *MUC4* alleles are closer the nuclear edge than the *LMNA*-WT and *LMNA*-MUT alleles.

