## Supplementary information for "Achieving single nucleotide sensitivity in direct hybridization genome imaging"

**Supplementary Table 1| Comparison of sgGOLDFISH with other nuclear DNA FISH methods.** The CasPLA method relies on Cas9's binding specificity to discriminate SNVs, therefore limits the target SNVs within a protospacer and proximal to (< 10 bp) the protospacer adjacent motif (PAM)[1]. sgGOLDFISH relies on eCas9 nickase's cleavage specificity to discriminate SNVs, hence allows for targeting SNVs distal to PAM. Moreover, existing methods visualizing SNVs in nuclear DNA involve proteinase treatment that excludes concurrent immunofluorescence and unavoidably perturbs nuclear architecture[1, 2]. In contrast, proteinase treatment is dispensable for sgGOLDFISH. Discriminating the *LMNA*-MUT allele from the *LMNA*-WT allele requires sgGOLDFISH to distinguish the G-U wobble base pair from A-U base pair (Extended Data Fig. 8c and 8d), which has been shown to be difficult using oligonucleotide probes alone such as in the Zombie method[3]. Also, Zombie is limited to detect SNVs in pre-integrated DNA barcodes because it requires phage promoters upstream of the target SNV[3], therefore it is not suitable for detecting endogenous nuclear SNVs.

| | **SNV-sensitive FISH methods** | | | | |
|---|---|---|---|---|---|
| | sgGOLDFISH | CasPLA[1] | Target-primed RCA of circularized padlock probes[4-6] | STAR-FISH[2] | Toehold probe strategy[7-9] |
| **Probe hybridization sites** | Target chromatin (1kb-1.5kb flanking target SNV) | *In situ* RCA amplification products (160kb-1Mb) | *In situ* RCA amplification products (160kb-1Mb) | *In situ* PCR amplification products (39 cycles of PCR) | Target RNA |
| **Target SNV location** | PAM-distal DNA (Yes) PAM-proximal DNA (possible) | PAM-proximal DNA | RNA | 3' end of a PCR primer | RNA |
| **Protein immunofluorescence compatible** | Yes | No | Possible | No | Possible |
| **Global DNA denaturation** | Not Required | Not Required | Not Required | Required | Not Required |
| **Labeling efficiency of non-repetitive targets** | ~25% | ~60% | 10%-30% | High | 50%-65% |

**Supplementary Note**

Sequencing-based methods have been developed for mapping single-strand breaks (SSBs) or double-strand breaks (DSBs) in cells, but they are expensive and do not provide absolute value of the fraction of DNA carries the breaks at a target site in a cell population[10-14]. A previously developed droplet digital PCR (ddPCR) assay unbiasedly measures the fraction of DNA with DSBs at a target site, but it is insensitive to SSBs. To quantitatively measure the cleavage efficiency of eCas9 nickase in complex with gMUC4-OneMM or gMUC4-twoMM in fixed cells, we modified the ddPCR assay to make it SSB-sensitive (therefore we call the modified ddPCR assay as SSB-ddPCR assay). The key modification is that we converted the SSB to DSB by an additional Cas9 nickase treatment.

In the Step 1 of the SSB-ddPCR assay, eCas9 nickase with gMUC4-OneMM or gMUC4-TwoMM was applied to fixed and permeabilized HEK293T cells to cleave its target genomic DNA, which would introduce a SSB if cleavage occurred (Extended Data Fig. 3a, Step 1, a SSB was introduced to the top strand of genomic DNA if cleavage occurred). The cells were then treated with proteinase K and the genomic DNA was harvested. In the Step 2, Cas9 nickase RNP (400 nM) which cleaves the other strand (i.e., the bottom strand in Extended Data Fig. 3a) was mixed with less than 8 ng/μL the harvested genomic DNA (Extended Data Fig. 3a, one or more SSBs were introduced to the bottom strand of genomic DNA in the Step 2 because multiple guide RNA species targeting the bottom strand were used to maximize cleavage efficiency). We found the efficiency of the Cas9 nickase RNP to cleave the bottom strand in the Step 2 was around 100% under our experimental condition (Extended Data Fig. 4a and 4b). Therefore, after the **eCas9** nickase RNP treatment in the Step 1 and the **Cas9** nickase RNP treatment in the Step 2, the target DNA was cleaved either on both strands or on the bottom strand only, depending on whether the eCas9 nickase RNP cleaved target DNA or not in the Step 1 (Extended Data Fig. 3a).

Next, the genomic DNA was purified to remove Cas9 nickase RNP and was mixed with two pairs of primers and two probes for ddPCR (Extended Data Fig. 3a). The F1/R1 primers span the cleavage sites of the eCas9 nickase RNP in the Step 1 and the Cas9 nickase RNP in the Step 2, while the F2/R2 primers do not (Extended Data Fig. 3a). The F1/R1 and F2/R2 amplicons are spaced by 216 base pairs. The amplification of the F1/R1 and F2/R2 amplicons are detected using FAM-quencher probe and HEX-quencher probe, respectively. In the ddPCR reactions, the DNA polymerase digests the probe annealed to template DNA by using its proofreading exonuclease activity, and releases the fluorescent dye from the quencher[15]. Therefore, a droplet shows FAM fluorescence or HEX fluorescence indicates amplification of the F1/R1 amplicon or the F2/R2 amplicon, respectively (Extended Data Fig. 3a). Hereinafter we call droplets with negative FAM signal and positive HEX signal as "- FAM + HEX droplets" (e.g., the green spots in Extended Data Fig. 3b), and the droplets with positive FAM signal and positive HEX signal as "+ FAM + HEX droplets" (e.g., the orange spots in Extended Data Fig. 3b). The fraction of "- FAM + HEX droplets" was calculated by using the number of "- FAM + HEX droplets" divided by the total number of "- FAM + HEX droplets" and "+ FAM + HEX droplets". When gMUC4-TwoMM, which contains two mismatches against the target DNA, or gMUC4-OneMM, which contains one mismatch, was complexed with eCas9 nickase for the SSB-ddPCR assay in HEK293T cells, we observed the fractions of "- FAM + HEX droplets" were 0.283 ± 0.005 and 0.580 ± 0.010, respectively (Extended Data Fig. 3b and 3c).

Ideally, if there is no DNA cleavage by eCas9 nickase RNP in the Step 1, the input DNA for ddPCR should have only a SSB on the bottom strand within the F1/R1 amplicon, and the ddPCR should generate "+ FAM + HEX droplets" because the top strand is intact (Extended Data Fig. 3a). However, even when we used catalytically dead Cas9 (dCas9)[16] instead of eCas9 nickase in the Step 1, the fractions of - FAM + HEX droplets was 0.294 ± 0.015 (Extended Data Fig. 4c). Similarly, in the previous study of DSB-ddPCR, about 7% droplets showed failed amplification of the F1/R1 amplicon and successful amplification of the F2/R2 amplicon even though the input DNA was uncleaved control DNA[15]. We speculate that the PCR amplification may fail in a fraction of droplets even though intact template DNA presents in the droplets, resulting in the "background level" of "- FAM + HEX droplets".

To obtain the absolute value of the fraction of DNA cleaved by eCas9 nickase RNP in the Step 1, we generated a standard curve that can convert the ddPCR readout (i.e., the fraction of "- FAM + HEX droplets") into the absolute value of the fraction of DNA cleaved in the Step 1 (Extended Data Fig. 3c). To generate the standard curve, dCas9 RNP (using gMUC4-OneMM) was applied to fixed and permeabilized cells, and then genomic DNA were harvested (Extended Data Fig. 5, Step 1). Next, the harvested genomic DNA was split into two tubes (Extended Data Fig. 5, Step 1). One was treated with restriction enzyme (Msel) to generate "ds-cut genomic DNA", and the other tube was treated with Cas9 nickase RNP which cleaves the bottom strand of target genomic DNA to generate "ss-nicked genomic DNA" (Extended Data Fig. 5, Step 2). Finally, "ds-cut genomic DNA" and "ss-nicked genomic DNA" were mixed at different ratios for ddPCR (Extended Data Fig. 5). The relationship between ddPCR readout and the fraction of "ds-cut genomic DNA" added is linear (Extended Data Fig. 3c, Pearson's $r^2 > 0.99$). According to the standard curve, the fraction of DNA cleaved by eCas9 nickase RNP in the Step 1 of the SSB-ddPCR was insignificant (< 0.05) for gMUC4-TwoMM and ~ 0.4 for gMUC4-OneMM (Extended Data Fig. 3c), consistent with the *in vitro* cleavage data using eCas9 RNP (Extended Data Fig. 2d).

**Supplementary information references**
